# Spinal V2a neurons support locomotor propagation via long-range connectivity

**DOI:** 10.64898/2026.09.23.753244

**Authors:** Saul Bello-Rojas, Chunqi Tian, Katarzyna M. Piekarz, Mary Niehaus, Martha W. Bagnall

**Author notes:** These authors contributed equally (Saul Bello-Rojas, Chunqi Tian).

## Abstract

The spinal cord contains a diverse set of neurons whose activity is vital for motor control. Despite the importance of these spinal interneurons in network models of locomotion, their circuitry remains poorly understood. We quantified the synaptic output of V2a (vsx2^+^) interneurons, a glutamatergic population implicated in control of locomotor speed, rhythm generation, and recovery from injury, using the larval zebrafish. Surprisingly, we found that the majority of monosynaptic output from V2a neurons occurs onto motor and premotor targets at long ranges, in contrast to models that posit V2a neurons as the source of recurrent local excitation. Focal loss of V2a neurons impairs long-range propagation of locomotor output, but not local rhythm generation, in accordance with their long-range connectivity. Our results position V2a neurons as major mediators of locomotor propagation and long-range coordination in the spinal cord.

## Introduction

The spinal cord is home to a wide diversity of interneurons whose collective activity is vital for motor and sensory functions. In the absence of descending or afferent input, the spinal cord can produce rhythmic, patterned motor output. This property is known as central pattern generation, and it has been intensively studied as it may explain key features of vertebrate locomotion and other rhythmic behaviors. Many spinal interneurons in the ventral portion of the cord participate in this rhythmic, patterned activity, and selective deletion or suppression of specific populations of spinal interneurons can alter the speed or pattern of motor output (Böhm et al., 2022; Callahan et al., 2019; Crone et al., 2008; Crone et al., 2009; Lanuza et al., 2004; Talpalar et al., 2013; Yao et al., 2025; Zagoraiou et al., 2009; Zhang et al., 2008). Accordingly, many models of spinal motor function rely on intricate patterns of connectivity among spinal interneurons (Cohen et al., 1992; Danner et al., 2017; Komi et al., 2024; Li et al., 2007; Roussel et al., 2021; Wallen et al., 1992; Zhang et al., 2022).

Despite the importance of these interneurons, their circuit interconnections are not well described, in part due to the difficulties associated with identifying and selectively stimulating these intermingled populations (Koch and Levine, 2023; Sengupta and Bagnall, 2023). A second challenge inherent to studies of the spinal circuit is that many interneurons project axons along the longitudinal (rostrocaudal) axis of the spinal cord, requiring analysis in the intact animal to preserve the spatial structure of these long-range connections. Thus, although there is a wealth of data analyzing local connectivity from interneurons onto motor neurons, we know far less about long-range connections and connectivity between interneuron populations. As a consequence, it has been a challenge to link abstract models of central pattern generator (CPG) circuitry with the actual neuronal populations present in spinal motor circuits.

V2a neurons have been of particular interest to the spinal motor field because of their influence on motor output and their proposed role as the rhythmogenic core of the CPG. V2a neurons form direct synaptic connections onto motor neurons and other V2a neurons, in both mice and fish (Ampatzis et al., 2014; Kimura et al., 2006; Song et al., 2020; Zhong et al., 2010). V2a neurons also share similar morphological subtypes across species, with some projecting purely descending and some bifurcating ascending/descending axons (Hayashi et al., 2018; Menelaou et al., 2014). Activation of V2a/vsx2^+^ spinal or brainstem neurons is sufficient to elicit locomotor output (Carbo-Tano et al., 2023; Cregg et al., 2020; Kimura et al., 2013; Ljunggren et al., 2014; Usseglio et al., 2020), while deletion or suppression of V2a spinal neurons impairs locomotor output (Crone *et al*., 2008; Eklof-Ljunggren et al., 2012). In conjunction with their connectivity, these results have placed V2a neurons as prime contenders for the role of rhythmogenic drivers of the fish CPG. In mouse, V2a neurons alone do not seem to be sufficient to explain the rhythmogenic drive, as deletion of V2a neurons leads to deficits in left-right coordination or decreases in frequency but not a total loss of rhythmogenesis (Crone *et al*., 2008; Crone *et al*., 2009; Yao *et al*., 2025). Instead, work points to a possible role for shox2^+^, hb9^+^, and/or spinocerebellar populations as important for rhythmic drive (Caldeira et al., 2017; Chalif et al., 2022; Dougherty et al., 2013; Ha and Dougherty, 2018). In fish, by contrast, the dominant model in the field identifies the recurrent excitatory drive from V2a neurons onto other local V2a and motor neurons as the source of rhythmogenic activity (El Manira, 2026; Grillner and El Manira, 2020; Grillner and Kozlov, 2021).

Given the importance of V2a neurons to many models of spinal motor control and efference copy (Azim et al., 2014; Roussel *et al*., 2021; Rybak et al., 2015), not to mention their potential role in recovery from injury (Jensen et al., 2024; Kathe et al., 2022), we sought to determine V2a output connectivity in a systematic, quantitative, spatially defined fashion along the long axis of the spinal cord. We applied optogenetic-based circuit mapping in the intact animal, as we have previously done to tackle longitudinal connectivity of other populations (Sengupta et al., 2025; Sengupta et al., 2021). Our results reveal that V2a neurons exhibit preferential long range, not local, outputs onto five spinal populations, including motor neurons, other V2a neurons, V1 neurons, and dI6 neurons. These results are at odds with a model in which highly local, recurrent V2a output is the primary source of rhythmogenesis. Instead, we find that selective V2a ablation impairs long-range, but not local, motor output. Our data indicate that V2a neurons mediate locomotor propagation and long-range coordination in the spinal cord. These results, in conjunction with prior work, present a new framework in which local inhibition and long-range excitation can produce rapid, robust movement of activity down the spinal cord.

## Results

### Optogenetic calibration of V2a spiking in zebrafish spinal cord

To create a map of V2a connectivity via optical stimulation, we generated a transgenic larval zebrafish line, Tg (*vsx2:Gal4:UAS:CatCh*), in which V2a neurons express the cationic channelrhodopsin CatCh (Fig. 1A). We designed and calibrated an optogenetic approach that would evoke minimal V2a spiking using a digital micromirror device (DMD). To validate the approach, we recorded evoked spiking from V2a neurons in whole-cell current clamp configuration during presentation of a 6x5 grid of localized optical stimuli over a single spinal muscle segment. This highly localized stimulation evoked spiking in V2a neurons only when the stimulus was presented on or adjacent to the soma (Fig. 1B, red traces). Elsewhere within the same segment, optical stimulation evoked subthreshold responses (black traces). 23/26 V2a neurons (88.5%) were driven to spike by this stimulus, validating the efficacy of this approach. Furthermore, stimulation outside the segment, 1-2 segments rostral or caudal to the recording site, failed to evoke spiking in this example neuron (Fig. 1C) and in 20/23 V2a neurons (Fig. 1D). Thus, the optical stimulus does not elicit antidromic spiking. We tested whether the absence of antidromic spiking was due to the small, low intensity stimulus by also presenting a full-segment, high-intensity optical stimulus. V2a neurons were optically activated when this stimulus was presented to the same segment as the cell body, as expected. However, this more intense stimulus could also elicit antidromic spiking when presented 1-2 segments caudal to the recording site (Supp. Fig. S1). Therefore, the spatially delimited stimulus is required to ensure that evoked spiking is confined to neurons with cell bodies in the illuminated segment. The absence of evoked spiking during out-of-segment stimulation with the small grid stimulus supports the contention that antidromic spiking, and polysynaptic transmission, are rare in this experimental configuration. For the remainder of the experiments, we used this small grid stimulus to activate V2a neurons sparsely, limiting the potential for polysynaptic transmission.

**Fig. 1.**
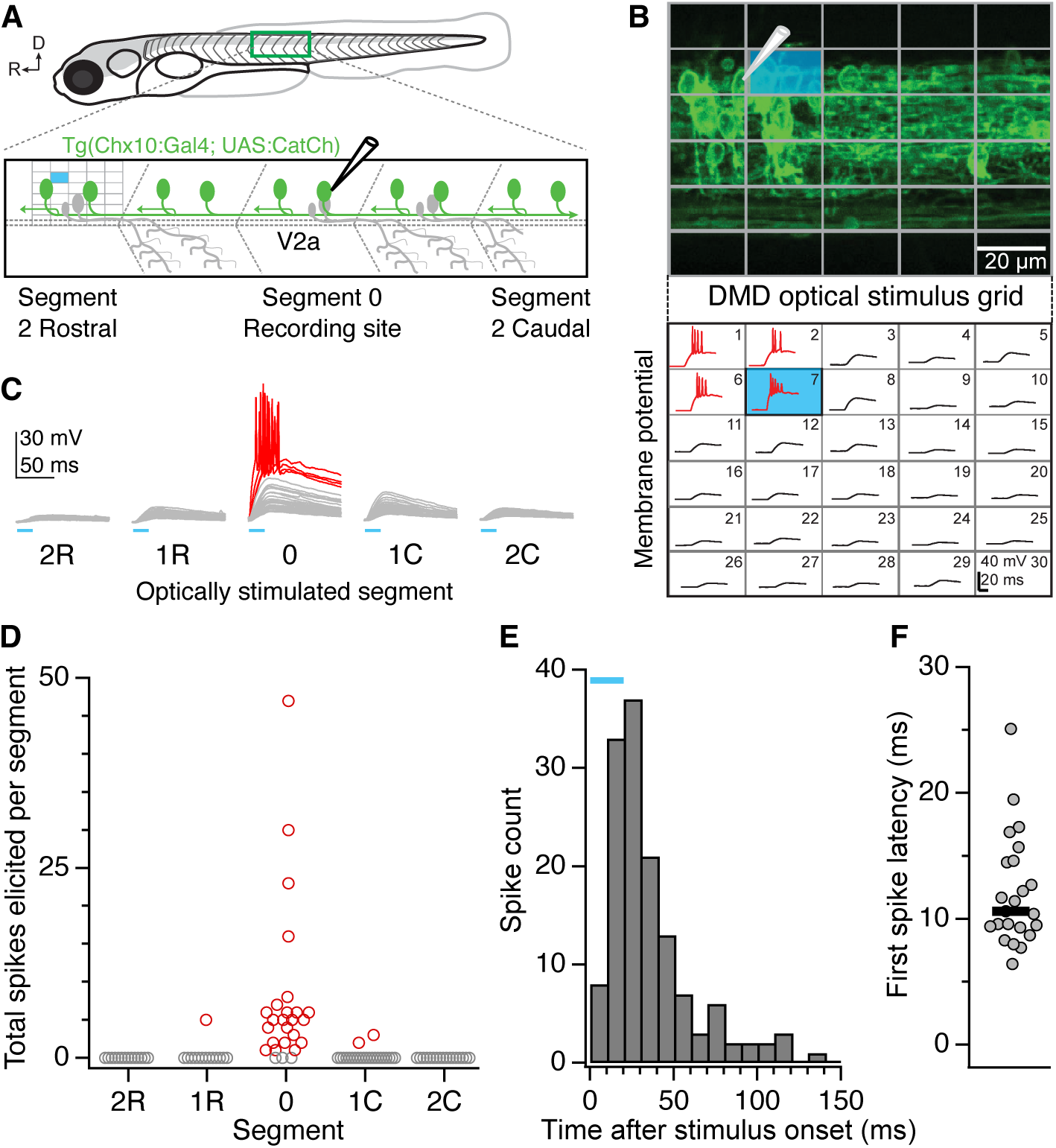
Calibration of optical stimulation of V2a neurons. **A,** Schematic of recording setup from V2a neurons in midbody spinal segments, with an example stimulus grid over segment 2R. **B,** Example segment image and V2a recording, showing spikes elicited by optical stimuli on or near the soma and its dendrites, but subthreshold responses elsewhere in the segment. **C,** Results from the same neuron during optical stimulation up to two segments rostral or caudal to the recording site. Only stimulation within the recorded segment elicits action potentials. **D,** Summary data as in C for 26 V2a neurons. Neurons with evoked spiking are shown in red. **E,** V2a neuron spiking relative to the duration of the optical stimulus (blue bar). **F,** First spike latency is typically ∼10 ms after onset of optical stimulus.

To determine the time period over which postsynaptic neurons would be likely to receive direct synaptic input from these optically activated V2a neurons, we measured the duration of elicited spiking. 96% of spikes were elicited within 100 ms after stimulus onset (Fig. 1E), and most optically activated V2a neurons initiated spiking within 20 ms of stimulus onset (first spike latency median 10.6 ms, Fig. 1F). We therefore used a 100 ms analysis window to detect optically evoked presumed monosynaptic EPSC events in our target neurons.

### V2a neurons excite motor neurons at long range

Motor neurons are a primary target of V2a neuron excitation, in both mice and zebrafish (Al-Mosawie et al., 2007; Ampatzis *et al*., 2014; Eklof-Ljunggren *et al*., 2012; Hayashi *et al*., 2018; Kimura *et al*., 2006; Menelaou and McLean, 2019; Stepien et al., 2010). However, little is known about the structure of V2a connectivity to motor neurons along the longitudinal axis of the spinal cord. Because V2a neurons make long, and in some cases bifurcating, projections along the rostrocaudal axis (Hayashi *et al*., 2018; Menelaou and McLean, 2019; Menelaou *et al*., 2014), it is possible that they connect to motor neurons broadly or in a more spatially delimited fashion. We systematically examined monosynaptic connectivity from V2a to motor neurons by recording from identified motor neurons while optically stimulating V2a neurons in different segments rostral or caudal to the recording site. The optical stimulus was delivered one segment at time, up to 6 segments rostral and caudal relative to the recording site (Fig. 2A). As the zebrafish comprises ∼30 muscle segments, this stimulation territory covers > 40% of the body. In this and subsequent experiments, target neurons were held at -80 mV in whole-cell voltage clamp configuration with a potassium-gluconate based internal solution to isolate EPSCs.

**Fig. 2.**
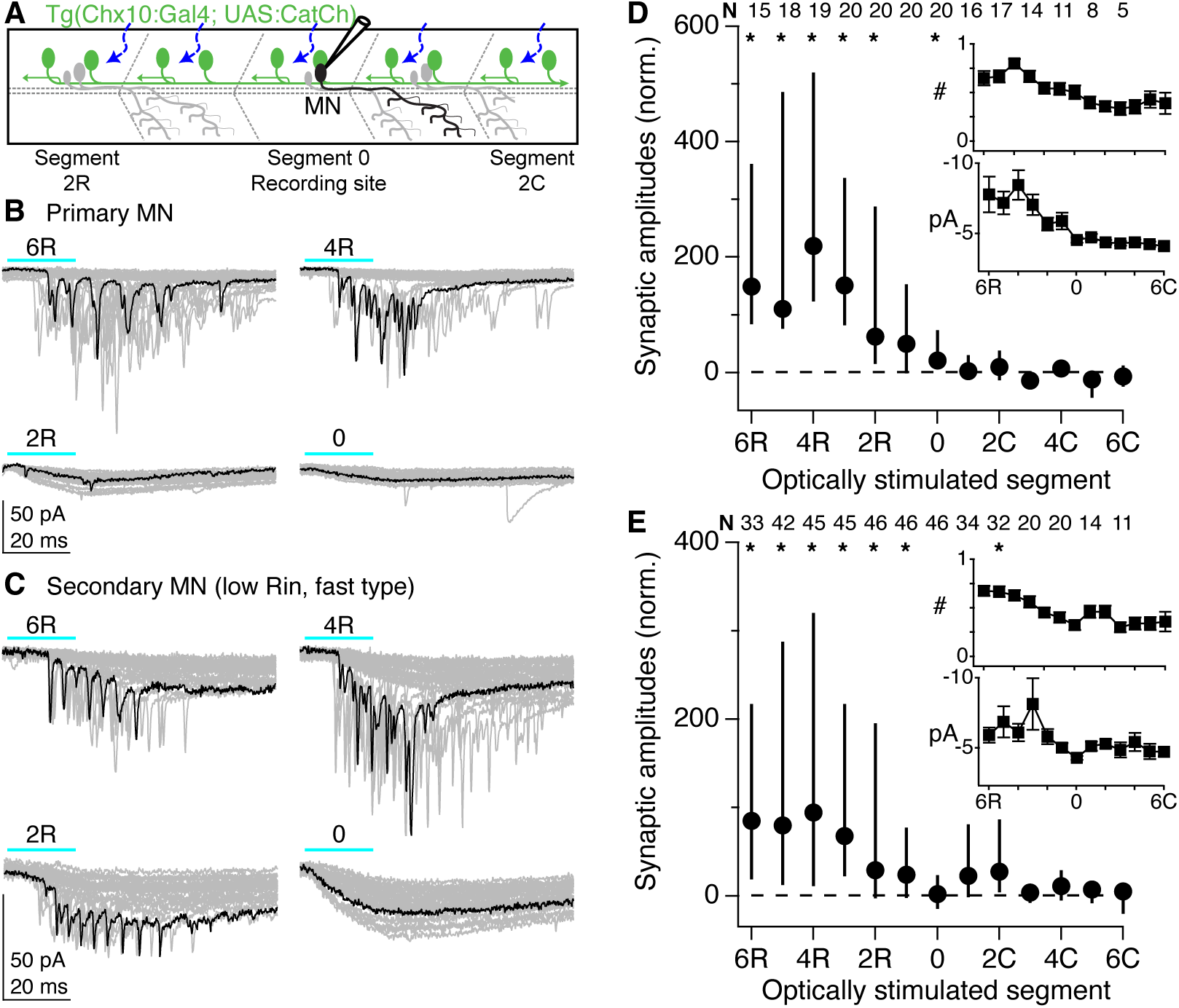
V2a neurons preferentially excite motor neurons at long range. **A,** Schematic of recording setup. Motor neurons were identified by axonal arborization into the muscle. Optical stimuli were delivered up to 6 segments rostral or caudal to the recording site. **B,** Example EPSCs recorded in a primary motor neuron. All the traces from one segment of stimulation are shown (gray), with a representative recording highlighted in black. **C,** As in **B** but for a fast secondary motor neuron. **D, E,** Summary data on the total sum of EPSC amplitudes, normalized to the recorded neuron’s intrinsic conductance, for primary and secondary motor neurons respectively. Data are shown as medians ± interquartile range. N = 20 primary and 48 secondary motor neurons; the number of neurons tested at each stimulation site is shown above the plot. Asterisks represent significant difference from baseline spontaneous events (Wilcoxon test), corrected for multiple comparisons. See Supp. Table S1. Inset top, number of EPSCs elicited by stimulation in each segment, normalized to the maximum number of EPSCs elicited per cell. Inset bottom, synaptic amplitudes (in pA) elicited by stimulation in each segment. Data are shown as means ± SEM.

Using post-recording cell fills, motor neurons were identified by their axonal arborization in the muscle. We further categorized cells as primary motor neurons (N = 20), fast secondary motor neurons (m- and ms-type, N = 34), or slow secondary motor neurons (s-type, N = 14) based on their soma size, input resistance, and axon arborization patterns, which are well characterized in the young zebrafish (Bello-Rojas et al., 2019; D’Elia et al., 2023; Menelaou and McLean, 2012). To our surprise, optical activation of V2a neurons locally to the recording site (between 2 segments rostral and 2 segments caudal to the recording site) evoked very few fast EPSCs in all classes of motor neurons. Instead, most motor neurons exhibited a slow inward current, with occasional small EPSCs with fast kinetics (Fig. 2B, C, segments 0, 2R). Optical stimulation 4-6 segments rostral to the recording site, in contrast, evoked barrages of large EPSCs with fast kinetics (Fig. 2B, segments 4R, 6R). To compare results across neurons, we quantified fast EPSC amplitudes within 100 ms after the onset of optical stimulation and normalized their sum to the intrinsic cellular conductance of each neuron (i.e., the summed EPSC amplitude was multiplied by the input resistance). Regardless of speed preference, motor neurons were maximally excited by V2a stimulation 3-6 segments rostral to the recording site, (Fig. 2D, E). In contrast, V2a stimulation within the recorded neuron segment evoked very little fast synaptic excitation, in any class of motor neurons. Differences in both the number of evoked EPSCs and their individual amplitudes contributed to the overall difference in synaptic strengths evoked from stimulation in long-range vs local sites (insets, Fig. 2D, E). Thus, both the number of synaptically connected neurons and their individual synaptic weights are likely components of the strong long-range connectivity. We attempted to measure connectivity from V2a neurons even more rostral to the recording site (7+ segments), but these stimuli frequently elicited fictive swimming (Ljunggren *et al*., 2014), preventing further quantification.

### V2a neurons excite other V2a neurons at long range

V2a neurons excite other V2a neurons through a mixture of electrical and chemical synapses (Ha and Dougherty, 2018; Menelaou and McLean, 2019; Song et al., 2018; Song *et al*., 2020; Zhong *et al*., 2010). Local connectivity within V2a neurons, assayed in slices from mouse spinal cord, is strongest in neonatal animals between pairs of neurons with shared properties (Ha and Dougherty, 2018; Zhong *et al*., 2010), and anatomical analyses support the presence of local connections in young adult animals (Hayashi *et al*., 2018). However, it has been challenging to address the strength and postsynaptic partner identity of long-range connections. Using the same strategy as for motor neurons, we recorded from 20 V2a neurons during optical stimulation of presynaptic V2a neurons along the longitudinal axis of the spinal cord (Fig. 3A). Measurement of V2a-V2a connectivity within the segment of recording was not possible due to the large direct depolarizing current evoked by opsin activation nearby. However, stimulation of local V2a neurons typically evoked small synaptic responses (Fig. 3B, segments 1R, 2R). In contrast, optical stimulation 4-6 segments rostral to the recording site evoked large amplitude EPSCs (Fig. 3B, segments 4R, 6R), similar to results in motor neurons. Data across neurons, normalized to postsynaptic conductance, reveals a maximal V2a – V2a connection by stimulation 6 segments rostral to the recording site (Fig. 3C). This effect was primarily driven by differences in number of evoked synaptic events, with a smaller contribution from differences in amplitude (insets).

**Fig. 3.**
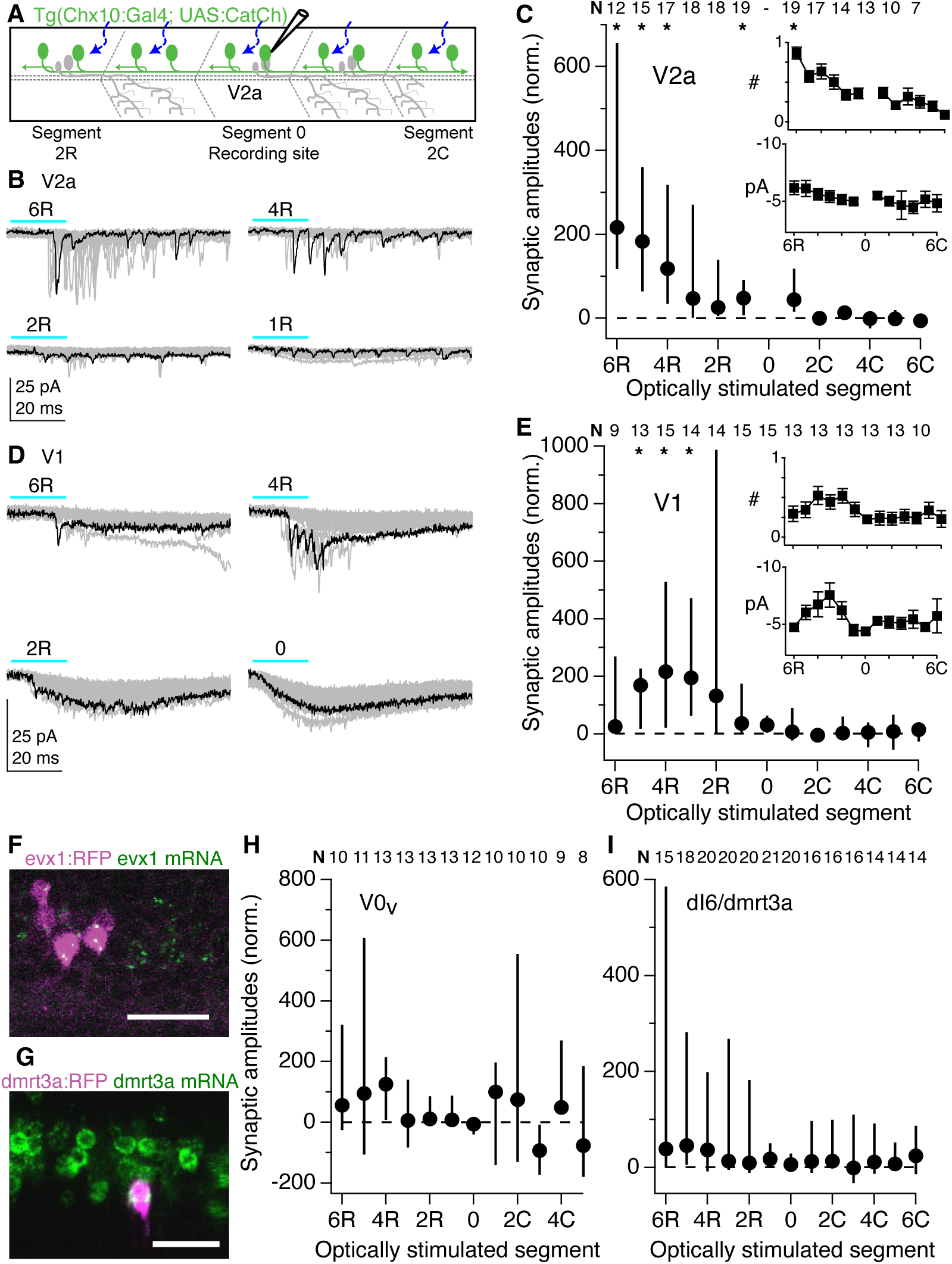
V2a neurons excite other ventral horn targets at medium-long range. **A,** Schematic of recording setup. **B,** EPSCs elicited in V2a neurons by optical stimulation of V2a neurons outside the recorded segment. **C,** Summary data for inputs onto V2a neurons, as in Fig. 2. **D, E,** as in **B, C** for V1 neurons. **F,** Image of HCR label of evx1 mRNA in the spinal cord of an animal expressing the evx1:RFP plasmid in a subset of neurons following embryo injection. RFP and mRNA signals colocalize, confirming the V0v identity of the RFP+ neuron. **G,** As in **F** but for dmrt3a mRNA and fluorophore label, confirming the identity of a subset of dI6 neurons. **H, I,** Summary of V2a inputs onto evx1+ V0v neurons (N = 13) and dmrt3a+ dI6 neurons (N = 21). Because of the modesty of the overall synaptic responses, EPSC numbers and amplitudes are not quantified.

### V2a neurons excite other spinal premotor neurons at medium-long range

V1 (En1^+^) neurons provide inhibition onto motor neurons, V2a neurons, and other premotor populations locally, while inhibiting sensory populations at long range (Li et al., 2004; Sengupta *et al*., 2021). V1 inhibition appears to help speed up locomotion by rapidly repolarizing motor pools, allowing them to cycle more quickly, in both mice and fish (Bhumbra et al., 2014; Gosgnach et al., 2006; Kimura and Higashijima, 2019). Computational modeling of the zebrafish axial circuit suggests that V2a neurons should excite V1 neurons (Roussel *et al*., 2021), but this prediction has not been examined in mice or zebrafish (Sengupta and Bagnall, 2023). We targeted V1 neurons for physiological recording in triple transgenic animals [Tg (*vsx2:Gal4;UAS:CatCh; en1:LRL:DTA*)]. As with other targets, optical stimulation of V2a neurons in the same segment as the recorded neuron evoked a slow inward current but few EPSCs (Fig. 3D). However, stimulation of V2a neurons 2-4 segments rostral to the recording site evoked large amplitude EPSCs, which tapered off in amplitude during stimulation in the most rostral segments (Fig. 3D). Across 20 V1 neurons, V2a-mediated excitation was maximal during stimulation 3-4 segments rostral to the recording site (Fig. 3E). Both the number and amplitude of evoked synaptic events tracked this pattern (insets). This distance represents somewhat closer range connectivity from V2a neurons onto V1 neurons than seen for motor or V2a targets, but still longer range connectivity than seen for V1 neurons onto V2a neurons (Sengupta *et al*., 2021).

In mouse, loss of V2a neurons produces deficits in left-right alternation, and anatomical evidence indicates synaptic connections from vsx2^+^ neurons onto commissural excitatory and inhibitory neurons (Crone *et al*., 2008). These connections also feature in network models of limbed locomotion (Danner *et al*., 2017). In fish, V2a neurons monosynaptically excite two classes of commissural inhibitory neurons (Guan et al., 2021; Menelaou and McLean, 2019) but the spatial extent is unknown, and there is no data on connectivity to commissural excitatory neurons. Therefore, we measured V2a synaptic connections onto two genetically defined populations of commissural neurons: the Evx1^+^ V0_V_ excitatory population, and the Dmrt3a^+^ dI6 inhibitory population. To visualize these populations, plasmids driving fluorescent reporters under control of the appropriate promoter were injected into *Tg (vsx2:Gal4;UAS:CatCh)* embryos at the single-cell stage for sparse expression. Construct labeling accuracy was validated with HCR in situ labeling. Dmrt3a mRNA was present in 95% of neurons fluorescently labeled following Dmrt3a:tdTomato plasmid injection (20/21 neurons in 9 fish), and Evx1 mRNA was present in 93% of neurons fluorescently labeled following Evx1:tdTomato plasmid injection (37/40 neurons in 11 fish) (Fig. 3F, G).

Optical stimulation of V2a neurons yielded variable synaptic responses in both commissural neuron populations, ranging from little to no response in some targets to large synaptic responses in others (Fig. 3H, I). The V0_V_ and dI6 populations as a whole received comparatively little V2a synaptic input overall, though some individual neurons received more than others. Collectively, V2a synaptic input onto these two genetically defined commissural populations appears modest, and as with other recorded targets, is weakest locally.

A summary of these connectivity patterns from V2a neurons onto different identified spinal targets across the rostrocaudal axis is shown in Fig. 4A. Here the data are displayed from the standpoint of the source V2a neurons, rather than from the standpoint of their recorded targets as in previous figures. Across the ventral horn, V2a neurons preferentially connect to their targets at medium to long range, with strongest connections 3-6 segments caudal, representing ∼10-20% of the body length. These results contrast with connectivity from ipsilaterally directed V1 and V2b inhibition, which is targeted to short-range ascending and short-to-medium range descending targets, respectively (Fig. 4B) (Bello-Rojas and Bagnall, 2022; Sengupta *et al*., 2025; Sengupta *et al*., 2021). These results suggest an overall structure of local inhibition and long-range excitation for the major ipsilaterally directed populations of the ventral horn.

**Fig. 4.**
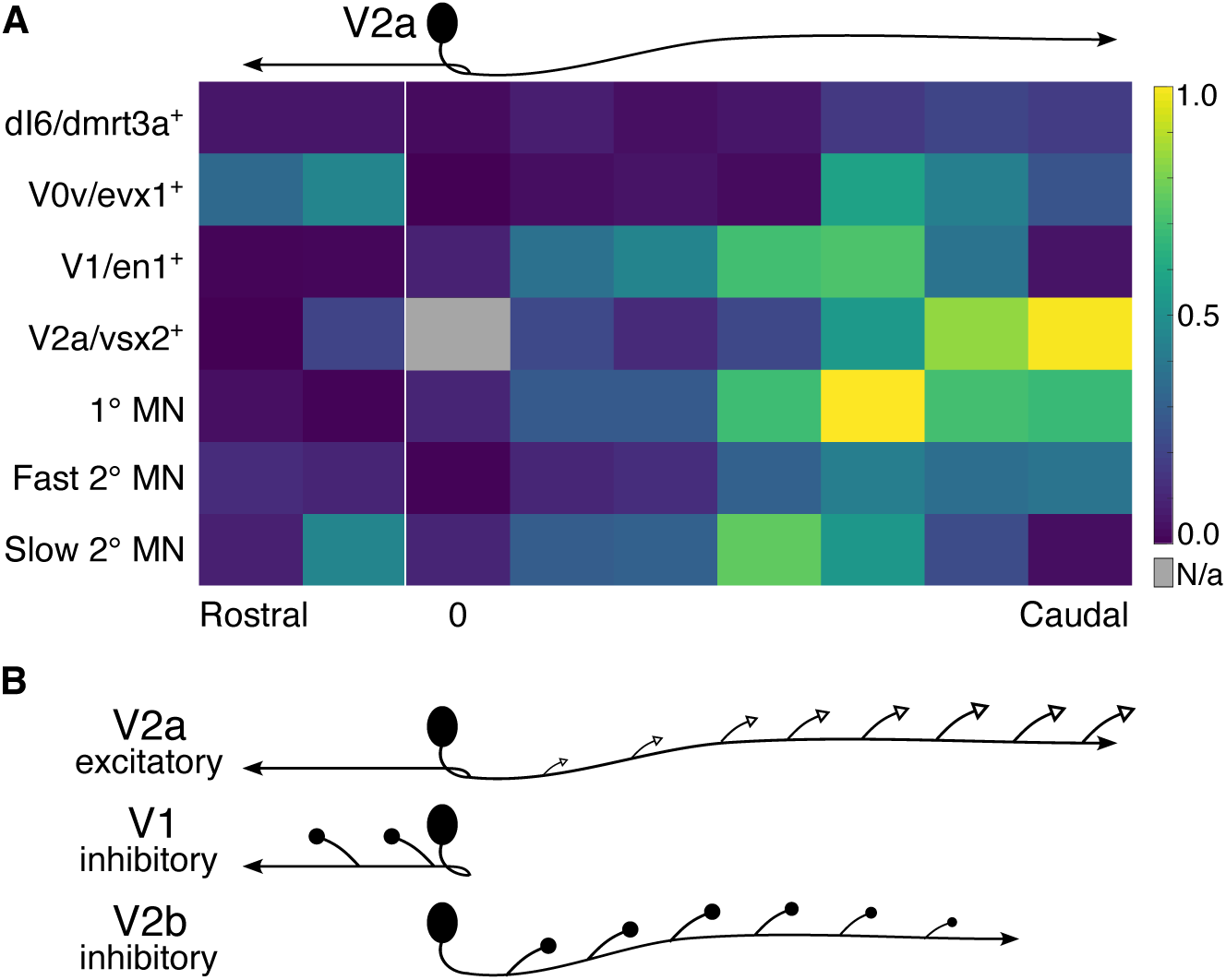
Summary of V2a connectivity along the rostrocaudal axis. **A,** Heatmap data representing normalized synaptic amplitudes for all targets as in previous figures, arranged relative to the standpoint of a presynaptic V2a neuron (that is, data from stimulation rostral to the recorded target neuron is displayed as caudal to the V2a neuron). Data from all recorded neuron target classes is normalized to the largest amplitude of all recordings to facilitate comparisons. Minimal local connectivity contrasts with strong long-range connectivity in the 3-6 segments caudal to the V2a neuron soma. **B,** Schematic of connectivity patterns for three classes of ipsilaterally projecting premotor spinal neurons. Stronger long-range excitation from V2a neurons (this paper) constrasts with stronger local and mid-range inhibition arising from V1 and V2b neurons (Sengupta et al., 2021; Sengupta et al., 2025; Bello-Rojas & Bagnall, 2022).

### V2a contributions to rhythm generation and propagation

This preferential long-range connectivity from V2a neurons onto ventral motor circuits appears at odds with the dominant view that V2a neurons provide local, recurrent excitation to support the key excitatory kernel (or “unit burst generator”) underlying rhythmic activity in the lamprey and zebrafish (Ampatzis *et al*., 2014; Grillner and El Manira, 2020; Grillner and Kozlov, 2021; Song *et al*., 2018). We reasoned that if V2a neurons are required for a local recurrent excitatory kernel underlying rhythmogenesis, then focal ablation in one region (3-4 segments) should be sufficient to impair motor activity in that area. In contrast, if V2a neurons subserve long-range functions, such as propagation of the locomotor wave down the spinal cord, then focal removal of V2a neurons should leave local motor generation unaffected, but impair downstream motor activity.

We first tested this reasoning in a 30-segment network model of the hemicord, using adaptive exponential integrate-and-fire neurons (Brette and Gerstner, 2005; Naud et al., 2008). Connectivity of inhibitory neurons was configured based on our published data, namely with V1 neurons providing short-range ascending inhibition and V2b neurons providing short- to medium-range descending inhibition (Supp. Fig. S3) (Sengupta *et al*., 2025; Sengupta *et al*., 2021). Connectivity of V2a neurons was defined by either our experimental measurements or a local recurrent architecture. In models with long-range V2a connectivity (Fig. 5A), as experimentally measured here, rostral drive initiated locomotor-like activity bouts characterized by brief burst firing and rapid propagation down the 30-segment model (Fig. 5B). Reducing output strength from V2a neurons located in midbody segments 9-12 to mimic ablation of part of the population caused a modest decrement in spiking (Fig. 5C). Across a range of tested reduced output strengths, these ablations still allowed for propagation of the activity burst (Fig. 5D). Motor neuron firing was minimally affected in the ablated segments, whereas downstream firing 5 segments caudal to the ablation was significantly reduced (Fig. 5E). In addition, ablation caused a delay in the onset of motor neuron bursts in caudal segments, slowing propagation speed and increasing intersegmental delay (Fig. 5F).

**Fig. 5.**
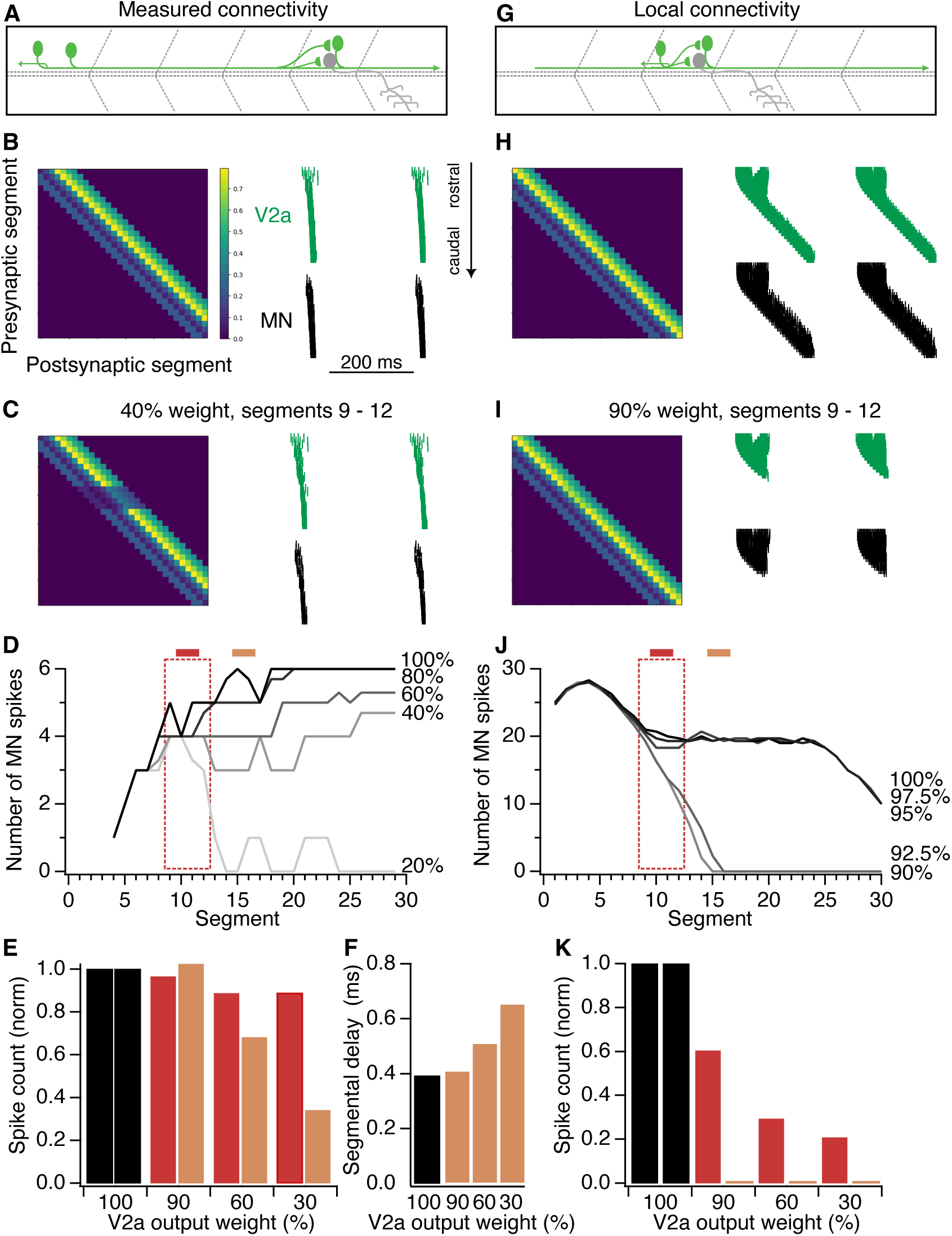
Two network models of V2a connectivity make different predictions. **A,** Schematic of V2a synaptic output as measured here with predominantly long-range connections. **B,** Weight matrix for V2a ◊ motor neurons based on data in Fig. 4 (left), and a raster of firing in V2a and motor neurons in a 30-segment model after initiating drive from rostral segments, showing activity propagation from rostral to caudal (right). **C,** Weight matrix after reducing output from V2a neurons located in segments 9-12 (left axis) onto their targets to 40% of original values. Activity continues to propagate to caudal segments, with a slight reduction in spiking. **D,** Summary of MN spiking in every segment under conditions of full strength connectivity (100%) or progressively reduced strength of synaptic output from V2a neurons located in the “ablated” segments (red dashed box). Activity propagates successfully, though with reduced motor neuron firing, until the weight is reduced to 20% of the original strength. **E,** Summary of motor neuron spiking in “ablated” segments (red) vs caudal to the ablation (brown), across several levels of weight reduction. **F,** Segmental delays in the segments caudal to the ablation increase with the degree of ablation. **G-K,** as for **A-E** but in a network with synaptic weights shifted in the rostrocaudal axis to model recurrent excitation. Despite much higher overall firing rates **(H)**, modest reductions in V2a synaptic output weight rapidly degrade performance, with activity unable to propagate at just 92.5% of the original weight matrix **(J)**. Segmental delays are not analyzed due to failures of propagation.

To compare this biologically inspired architecture with a locally recurrent network, we shifted synaptic weights in the rostrocaudal axis so that the peak synaptic output from a given V2a neuron onto motor neurons and other V2a neurons occurred within the same segment as its soma, but with otherwise identical weight distributions (Fig. 5G). In this model, locomotor activity was strong, with tens of spikes per motor neurons, but took longer to propagate down the 30 segment model (Fig. 5H). Despite the far larger number of spikes in each segment of this recurrent architecture, it was highly susceptible to loss of V2a synaptic output. Loss of 10% of V2a output strength in segments 9-12 was sufficient to completely abrogate locomotor propagation (Fig. 5I, J). Comparable ablations as those tested in the previous model yielded dramatically reduced motor neuron firing both in the ablated region and the region just caudal to the ablation (Fig. 5K). Due to the rapid degradation of performance, we could not explore effects of V2a ablation on propagation speed (cf. Fig. 5F). This greater susceptibility to ablation, despite identical model parameters and overall synaptic weights, occurs because the loss of local recurrent excitation rapidly compounds upon itself. Similar results were obtained in networks with ablations in a more caudal region (segments 17-20), demonstrating that these results were not tied to the differences in rostral firing patterns between the two models (Supplemental Fig. S4). Thus, if V2a connectivity is primarily long-range, then V2a ablation in a focal region is predicted to have minimal local effects and instead incur consequences on downstream activity rates. Conversely, if V2a connectivity is primarily local, ablations are predicted to impair both local and long-range motor output (cf. Figs. 5e, k).

We next tested these predictions experimentally. We used pulsed-laser ablation of V2a neurons in a 3-4 segment midbody region of agarose-embedded TgBAC(vsx2:EGFP)^nns1Tg^ animals, followed by 24 hr free-swimming to permit recovery from acute cell death-associated effects. To compare local vs long-range consequences of V2a ablation, we measured motor neuron activity via ventral root recordings, one within the ablated territory, and one in intact territory, 4-9 segments caudal to the first recording (Fig. 6A; median, 5 segments caudal). Ablation eliminated an average of 75 V2a neurons across 3-4 segments, or 70.7% ± 2.1% of the neurons present (mean ± SEM; n = 21). Animals that were agarose-embedded but not ablated served as controls.

**Fig. 6.**
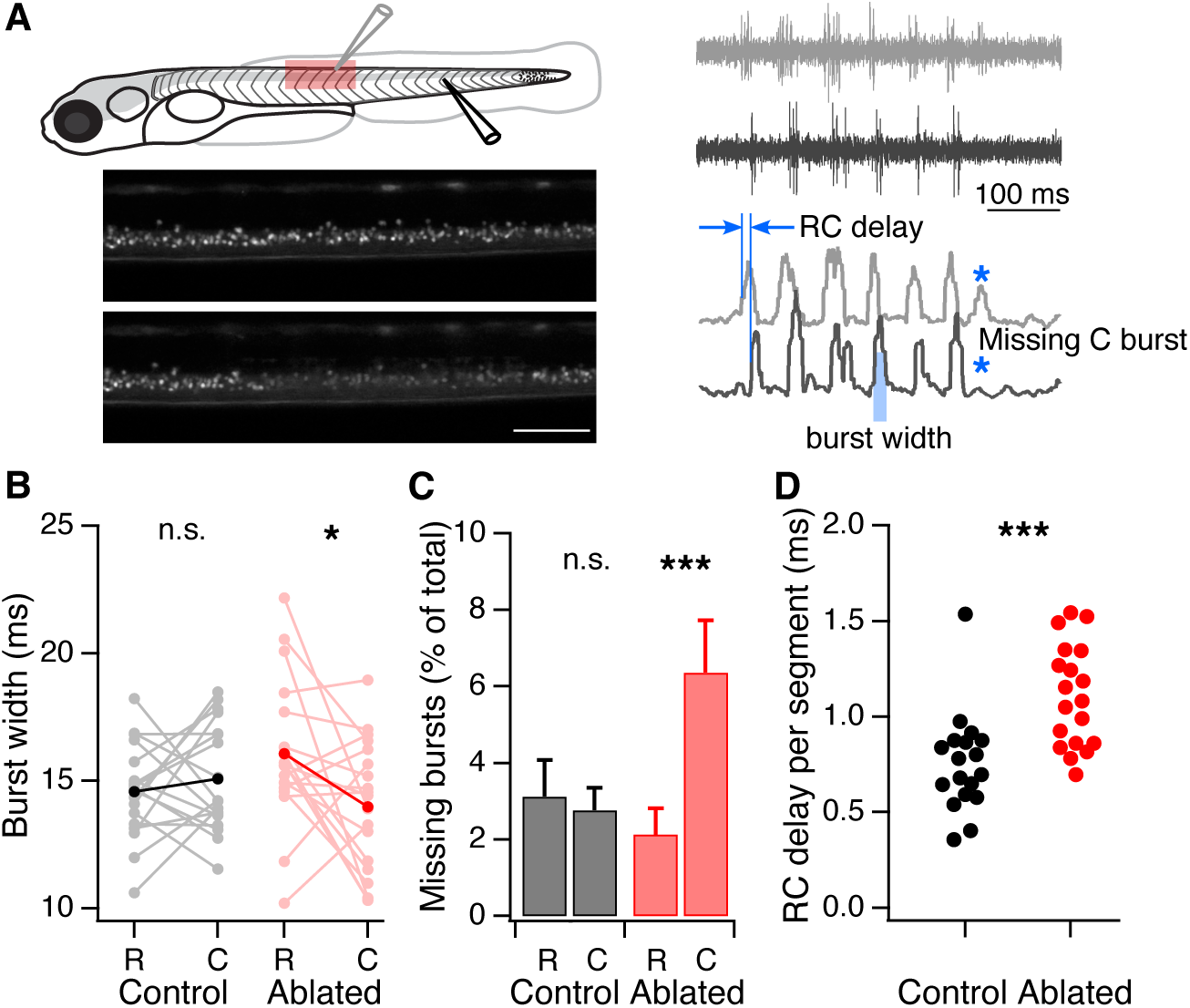
V2a ablation impairs propagation without affecting local drive. A, Schematic of preparation. V2a neurons were ablated over a 3-4 segment region at midbody; example before and after images, below (scale bar, 100 µm). Subsequent recordings of motor output during fictive locomotion were made simultaneously in the ablated region (rostral electrode, gray trace) and an intact region (caudal electrode, black). Smoothed standard deviation traces, below, were used for analysis. **B,** Motor power, measured as burst width, is weaker in the caudal than the rostral recording site, selectively in the V2a ablated animals. *, *p* = 0.029. **C**, The caudal but not rostral recording site exhibits elevated rates of “missing” bursts, indicating that rostrocaudal propagation is impaired in V2a ablated animals. ***, *p* = 0.0009. **D,** Onset of burst times occurs later in ablated animals, indicating that rostrocaudal propagation is slowed during locomotion. ***, *p* = 0.0005.

During fictive locomotion, motor neuron bursts were measurable in both the rostral (V2a ablated) segments and the caudal unablated segments, demonstrating successful propagation of swimming pattern past the ablated region (Fig. 6A). Ablation of V2a neurons did not affect the strength of the motor burst in the ablated region itself, but instead reduced burst width in the downstream caudal region (Fig. 6B). Because burst width is a proxy for motor neuron spiking, these results suggest a loss of motor neuron firing caudal to the ablation, but not within the ablated region, consistent with predictions from the network model endowed with experimentally measured connectivity. Similarly, we identified a number of locomotor bursts that occurred only in the rostral segment, without a matched caudal burst (example, Fig. 6A). Though ventral root recordings may not reliably capture all motor activity, these failures to propagate occurred at far higher rates in V2a ablated animals, compared to controls (Fig. 6C). Failures to propagate were most common in the last burst of a swim bout (77% of failures to propagate). In addition, rostrocaudal propagation times, measured from the onset of the rostral burst to the onset of the caudal one, were significantly slowed in V2a ablated animals (Fig. 6D). These results are consistent with a model in which motor activity is driven primarily by long-range, rather than local, V2a excitation.

## Discussion

Spinal V2a neurons are a major excitatory driver population, implicated in left-right alternation, rhythmogenesis, and recovery from spinal cord injury. By systematic mapping of their connectivity along the long axis of the spinal cord, we find that V2a neurons predominantly excite motor neurons and other ventral horn populations at long ranges, with relatively weak local synapses. Recordings of locomotor output demonstrate that loss of V2a neurons has little effect on local rhythmogenesis or motor activity, and instead impairs propagation and coordination between rostral and caudal sites. These results support a model in which V2a excitatory drive, in concert with local inhibitory circuits, effectively pushes a propagating wave down the spinal cord to support locomotion.

Connectivity motifs among populations of spinal interneurons are as yet poorly defined (Sengupta and Bagnall, 2023). Our work, in conjunction with earlier longitudinal mapping of spinal connectivity (Sengupta *et al*., 2025; Sengupta *et al*., 2021), suggests a basic structure of ipsilateral connectivity: V1 and V2b-mediated inhibition dominates locally, while V2a-mediated excitation builds up at longer range. This structure may provide clues to its functional operation. Local inhibition and long-range excitation are proposed to underlie temporally precise sequencing of activity in birdsong control circuits and entorhinal cortex (Kornfeld et al., 2017; Schmidt et al., 2017). Neural spaces with the opposite structure—local excitation flanked by long-range inhibition, such as in the fly heading direction complex (Green et al., 2017; Kim et al., 2017)—are thought to stabilize a bump of activity, which can then be modulated by inputs. The job of the spinal cord, in contrast, is to propagate activity, in both fish and tetrapods (Bonnot et al., 2002; Cuellar et al., 2009; Roussel *et al*., 2021). The circuit structure described here is inherently destabilized, supporting rapid propagation of a locomotor bend from head to tail.

Our connectivity map in young zebrafish stands in contrast to prior work in adult zebrafish demonstrating strong local connectivity among V2a neurons and from V2a to motor neurons (Ampatzis *et al*., 2014; Song *et al*., 2020). The discrepancy may be explained by animal age, as some features of the spinal network shift over development (Pallucchi et al., 2022; Picton et al., 2021). Another possible explanation is that the measured local connectivity in adult fish represents indirect electrical connectivity, rather than monosynaptic direct synaptic connections (Menelaou and McLean, 2019). A network in which V2a neurons are indirectly connected with other populations via gap junctional connections with a shared intermediate, such as a descending axon, is consistent with these observations. Support for this hypothesis comes both from prior circuit analysis as well as our own measurements of synaptic kinetics. Descending neurons make electrical synaptic connections onto V2a neurons and other targets, including motor neurons, in both larval and adult zebrafish (Guan *et al*., 2021; Pujala and Koyama, 2019; Wang and McLean, 2014), and indeed long-range V2a axons themselves synapse with both V2a and motor neurons (Figs. 2, 3). Small synaptic amplitudes and apparent slow rise times of local connections measured in adult animals (Ampatzis *et al*., 2014; Song *et al*., 2020) contrast with larger amplitudes and much faster (1-2 orders of magnitude) rise times of V2a synaptic connections in larval animals (Supp. Fig S2), supporting this hypothesis. Indeed, strong V2a synapses onto motor neurons occur at long distances in adult fish (Pallucchi *et al*., 2022), raising the possibility that this structure is retained in adult animals. However, systematic mapping or connectomic experiments along the long axis of the cord in adults will be required to determine the answer.

Rhythm generation by neurobiological networks has been a topic of intense interest. While many theoretical models can generate rhythmic oscillations, in relatively few experimental models do we have a mechanistic explanation (Marder and Bucher, 2001). Many circuit models of the spinal cord postulate a rhythmogenic kernel but do not identify a biological source (Danner *et al*., 2017; Yao *et al*., 2025). Recurrent excitation, paired with intrinsic physiological fatigue, has been proposed to underlie the spinal CPG, based on early observations of local excitatory connections (Buchanan and Grillner, 1987). The V2a population, as ipsilateral excitatory neurons rhythmically active during locomotion, has been a prime candidate to provide this hypothesized drive. In mouse, however, loss of V2a neurons does not abrogate rhythm generation (Crone *et al*., 2008; Yao *et al*., 2025). Other populations, such as Shox2^+^ or spinocerebellar neurons, have been suggested to fill the role (Chalif *et al*., 2022; Dougherty *et al*., 2013; Ha and Dougherty, 2018; Singh et al., 2025). Alternatively, inhibitory neurons can generate rhythms by enforcing alternation (Moult et al., 2013; Wandler et al., 2025), or rhythm may emerge from dynamic network activity (Linden et al., 2022). Our data demonstrating persistent, intact motor bursts arising from V2a-ablated segments of spinal cord do not support the idea that V2a neurons supply the drive for local motor burst generation. Instead, the decrease in downstream motor burst duration indicates that in larval zebrafish, V2a neurons carry a propagating wave of activity down the cord.

## Limitations of the study

While we sampled as comprehensively as we could from several spinal populations, some classes of ventral horn neurons have not been sufficiently assayed as possible targets of V2a excitation. These include the inhibitory V2b neurons and excitatory V3 neurons, as well as under-described populations like the sox1^+^ V2c/V2s (Gerber et al., 2019; Panayi et al., 2010). These or other ventral populations could be targets of local V2a output, which seems likely to exist based on anatomical analysis (Menelaou *et al*., 2014). One limitation of the optogenetic stimulation method is that, although optical stimuli are carefully calibrated, not all neurons may be equally well driven to spike, leading to uneven mapping results. Finally, all recordings were carried out in the midbody range for consistency; the structure of circuits may differ in far rostral or caudal positions. Therefore, connectivity onto untested targets, or from a low-excitability subset of V2a neurons, may be different than what is described here. Nonetheless, the consistency of results between connectivity mapping and behavior support the idea that most V2a drive onto motor circuits operates over long ranges.

## Materials & Methods

### Animal care and transgenic lines

All fish used for experiments were at the larval stage from 4 – 6 dpf, before onset of sexual differentiation. All experiments and procedures were approved by the Animal Studies Committee at Washington University and adhere to the NIH guidelines. Adult zebrafish (*Danio rerio*) were maintained at 28.5°C with 14:10 light:dark cycle in the Washington University Zebrafish Facility following standard care procedures. Larval zebrafish used for experiments were kept in Petri dishes in system water or housed with system water flow. Transgenic animals *Tg*(*vsx2:Gal4*;*UAS:CatCh*) were created by crossing the transgenic line *Tg*(*vsx2:Gal4;UAS:GFP*) (ZDB-FISH-220828-1) with a stable *Tg*(*UAS:CatCh*) line. For targeting V1 neurons, *Tg*(*eng1b:loxP-DsRed-loxP-DTA*) (ZDB-ALT-191030-2) were crossed to *Tg*(*vsx2:Gal4;UAS:CatCh*) to generate *Tg*(*vsx2:Gal4;UAS:CatCh; eng1b:loxP-DsRed-loxP-GFP*).

### V0v and dI6 identification

Embryos were injected with either a *de novo* generated *dmrt3a*:*mCherry* BAC or *evx1*:*mCherry* plasmid (VectorBuilder) at 10 – 15 ng/µl during the 1-2 cell stage (Juárez-Morales et al., 2016; Satou et al., 2020). At 4 dpf, larvae were screened for sparse expression of mCherry in the spinal cord and selected for electrophysiology or fluorescent *in situ* experiments.

Hybridization chain reaction (HCR) fluorescent *in situs* were performed in WT larvae injected with either *dmrt3a*:*mCherry* or *evx1*:*mCherry* plasmid to validate neuronal identity of stochastically labeled neurons. Synthesis of RNA probes designed for *dmrt3a* and *evx1* were performed as previously described (Thisse & Thisse, 2008; Tsai et al., 2020). We followed the protocols suggested by Molecular Instruments and adapted for 4 – 5 dpf larval zebrafish (Tsai et al., 2020). Hybridized larvae were mounted on slides using Vectashield (ThermoFisher) and cured for > 24 hr. Larvae were then imaged using an Olympus FV1200 Confocal microscope and a UAPO 40x W/340 1.15 NA water immersion objective. A transmitted light image was obtained along with laser scanning fluorescent images to identify spinal segments. Sequential scanning was used for multi-wavelength images. Confocal image stacks were reconstructed and analyzed using Imaris (9.8, Bitplane). Co-localization of native fluorescence and fluorescent *in situ* label was analyzed manually using the Slice function of Imaris. Co-localization frequency was calculated by taking the percentage of neurons that co-expressed the stated marker out of all neurons with reporter expression.

### Electrophysiology acquisition and analysis

Whole-cell patch-clamp recordings were performed in larvae at 4 – 6 dpf. Larvae were immobilized with 0.1% α – bungarotoxin and fixed to a Sylgard lined Petri dish with custom-sharpened tungsten pins. One muscle segment overlaying the spinal cord was removed at the mid – body level (segments 8 – 16) using a blunt-end glass electrode and suction (Wen and Brehm, 2010). The larva was then transferred to a microscope (Scientifica SliceScope Pro) equipped with infrared differential interface contrast optics, epifluorescence, and immersion objectives (Olympus: 40x, 0.8 NA). The bath solution consisted of (in mM) 134 NaCl, 2.9 KCl, 2.5 MgCl_2_, 10 HEPES, 10 glucose, and 2.8 CaCl_2_. The modestly high divalent concentration is helpful in reducing polysynaptic activity. Osmolarity was adjusted to ∼290 mOsm and pH to 7.5. Patch pipettes (5–15 MΩ) were filled with internal solution for voltage and current clamp composed of (in mM) 125 K gluconate, 2 MgCl_2_, 4 KCl, 10 HEPES, 10 EGTA, and 4 Na_2_ATP. Additionally, Alexa Fluor 647 hydrazide 0.05–0.1 mM or sulforhodamine (0.02%) was included to visualize morphology of recorded cells post hoc. Osmolarity was adjusted to ∼290 mOsm and KOH was used to bring the pH to 7.5. Patch recordings were made in whole-cell configuration using a Multiclamp 700B, filtered at 10 kHz (current clamp) or 2 kHz (voltage clamp). All recordings were digitized at 50 kHz with a Digidata 1440 (Molecular Devices) and acquired with pClamp 10 (Molecular Devices).

A Polygon 400 Digital Micromirror Device (Mightex) was used to deliver optical stimulation. The projected optical pattern consisted of a 6x5 grid of 30 squares. Each square in the grid measured ∼20 x 12 µm. For each square, illumination consisted of a 20 ms light pulse (470 nm) at 25% intensity (∼4 µW measured with 40X, 0.8 NA objective). The objective was positioned over a single spinal segment for stimulus delivery; the stage was translated and repositioned for each subsequent segment stimulated. V2a spiking was measured in current-clamp mode; synaptic inputs onto target neurons were measured in voltage-clamp mode (-80 mV), except for data in Fig. S2, obtained in current-clamp mode. The same protocol was used for all cell types and in all segments to obtain reliable EPSCs for measurement of synaptic input. For full-segment, high-intensity V2a stimulation, the laser intensity was increased to 100%, and the optogenetic pulse was presented to the whole segment being stimulated.

Electrophysiology data were imported into Igor Pro 9 (Wavemetrics) with Neuromatic (Rothman and Silver, 2018). Spikes and EPSCs were detected and analyzed using custom code in Igor Pro and MATLAB. The detection algorithm was based on the event detection instantiated in SpAcAN environment for Igor Pro (Bagnall and McLean, 2014; Dugue et al., 2005; Rousseau et al., 2012). Summed synaptic input for evoked responses was calculated by summing all detected EPSC event amplitudes within 100 ms of stimulus onset, and subtracting detected EPSC events within a 100 ms baseline period prior to the stimulus to account for spontaneous activity. All summed synaptic input values were then normalized by the total conductance (inverse of input resistance) of the neuron. Statistical comparisons were made between summed evoked EPSC amplitudes and summed spontaneous EPSCs occurring in a baseline period.

### V2a ablation and locomotor activity

4 dpf TgBAC(vsx2:EGFP)^nns1Tg^ larvae were anesthetized in 0.02% MS-222 and embedded in low-melting-point agarose (1%) in a 10mm FluoroDish (WPI). Targeted laser ablations were then performed using a Bruker Ultima 2pPlus two-photon microscope with a Coherent Monaco laser system. A 25x water-immersive objective (NA 1.1, Nikon) was used for ablations. Individual V2a cells were ablated by manually positioning the laser focus over the GFP-positive somata and delivering pulses at ∼37 mW at a wavelength of 1040 nm for 0.1 ms. Ablations were performed across approximately 4 spinal segments slightly rostral to midbody (∼segments 10-13) in each fish. After ablations, larvae were unembedded from the agarose and recovered for ∼24 hours in system water before post-ablation experiments. Control animals were similarly handled but not subjected to ablation.

To record the propagation of motor activity, animals were paralyzed and pinned down as above, and the skin was removed over segments ∼8-20. Dual ventral root recordings were performed using suction electrodes. The rostral electrode was positioned between segments ∼10-13, i.e. in one of the V2a-ablated segments or similar rostrocaudal position in control animals. The second electrode was positioned 4-9 segments caudal to the first electrode. Motor activity was then recorded during fictive locomotion, either spontaneous or evoked by full-field white light illumination. Signals were recorded in 100x AC mode, filtered to 300 Hz - 2 kHz, and digitized at 50 kHz. Fictive locomotion was quantified using the standard deviation of the ventral root recording over a 10 ms sliding window (Huang et al., 2013), thresholded to find events. Burst times were calculated as event midpoints. Events identified in one recording but not the other were visually examined to check for possible subthreshold activity.

### Network modeling

The simulation of the 30-segment zebrafish hemicord was coded in Brian2 (Stimberg et al., 2019), using an adaptive exponential leaky integrate-and-fire (AdEx) neuronal model (Brette and Gerstner, 2005; Naud *et al*., 2008). The model features four neuronal populations: V2a, MN, V1, V2b; each segment contains one neuron of each kind. The initial membrane parameters were based on the literature or from electrophysiology data collected by the laboratory (Buss et al., 2003; Menelaou and McLean, 2019). AdEx parameters were fitted for a single neuron of each population, using extensive grid searches, to achieve firing patterns that follow the experimental traces, and the neurons were then incorporated into a network containing 30 segments. Connectivity between the neuronal populations was based on our experimental data, and extended rostrally and caudally to cover all 30 segments (Supp. Fig. S2). AdEx parameters and synaptic connection strengths were adjusted to ensure full propagation in the experimentally-based model; the same parameters and synaptic weights were used in the locally connected, recurrent model. The six most caudal V2a cells receive a graded input current that jump-starts the rostrocaudal cycle. This input current is injected three times for 60 ms, which results in three cycles of activity within each simulation run.

### Statistics

Statistical tests were performed using MATLAB (R2018a, MathWorks) or Igor Pro 9. Due to the non-normal distribution of physiological results, nonparametric statistics and tests were used for representations and comparisons. Details of statistical tests, p-values, and sample size are described in the corresponding figures and legends, or in Supplemental Table 1.

## Author contributions

S. B.-R. and C.T. carried out and analyzed physiological and behavioral experiments. K.P. carried out computational modeling. M.N. carried out and quantified anatomical experiments. M.W.B. supervised all work, and wrote the manuscript with contributions from all authors.

## Acknowledgments

This work was supported by the National Institutes of Health (R01NS130483 to M.W.B.) and the McDonnell Center for Systems Neuroscience (M.W.B.). Ablations and imaging were performed in part through the use of Washington University Center for Cellular Imaging supported by Washington University School of Medicine, The Children’s Discovery Institute of Washington University and St. Louis Children’s Hospital (CDI-CORE-2015-505 and CDI-CORE-2019-813) and the Foundation for Barnes-Jewish Hospital (3770 and 4642).

## Materials availability

Transgenic lines and plasmids used in this study are available from the lead contact upon request.

