## Supplemental Figures for "Spinal V2a neurons support locomotor propagation via long-range connectivity"

**Supplemental figures and table**

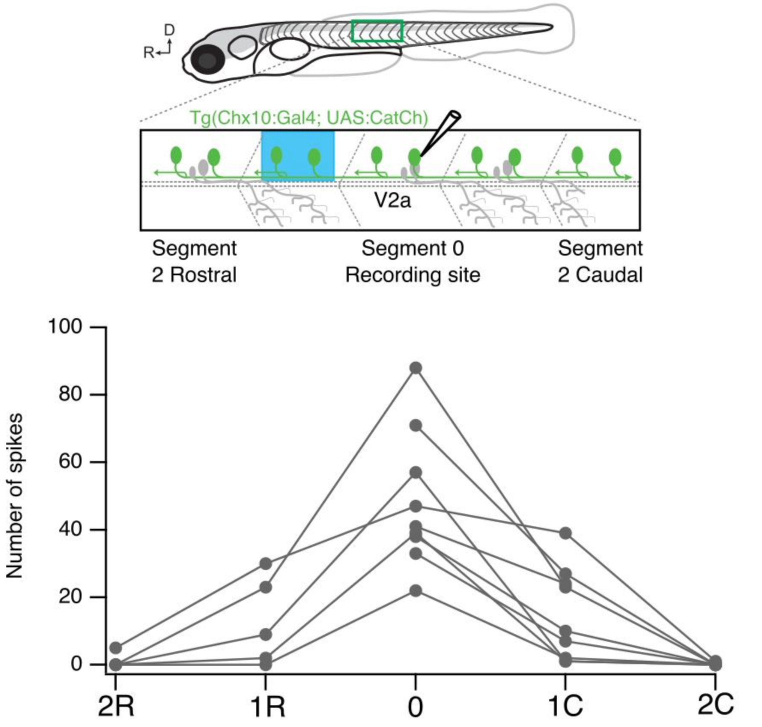


**Supp. Figure S1:** Full-segment optogenetic stimulation. As in Fig. 1, but the stimuli delivered by DMD covered the entire segment area rather than 1 of 30 grid squares within that area. Stimuli were also delivered at 100% stimulus strength (cf. 25% for main results). Under these conditions, optical stimuli delivered rostral or caudal to the recorded segment can often evoke spiking, presumably through antidromic activation and/or synaptic stimulation. N = 9 neurons.

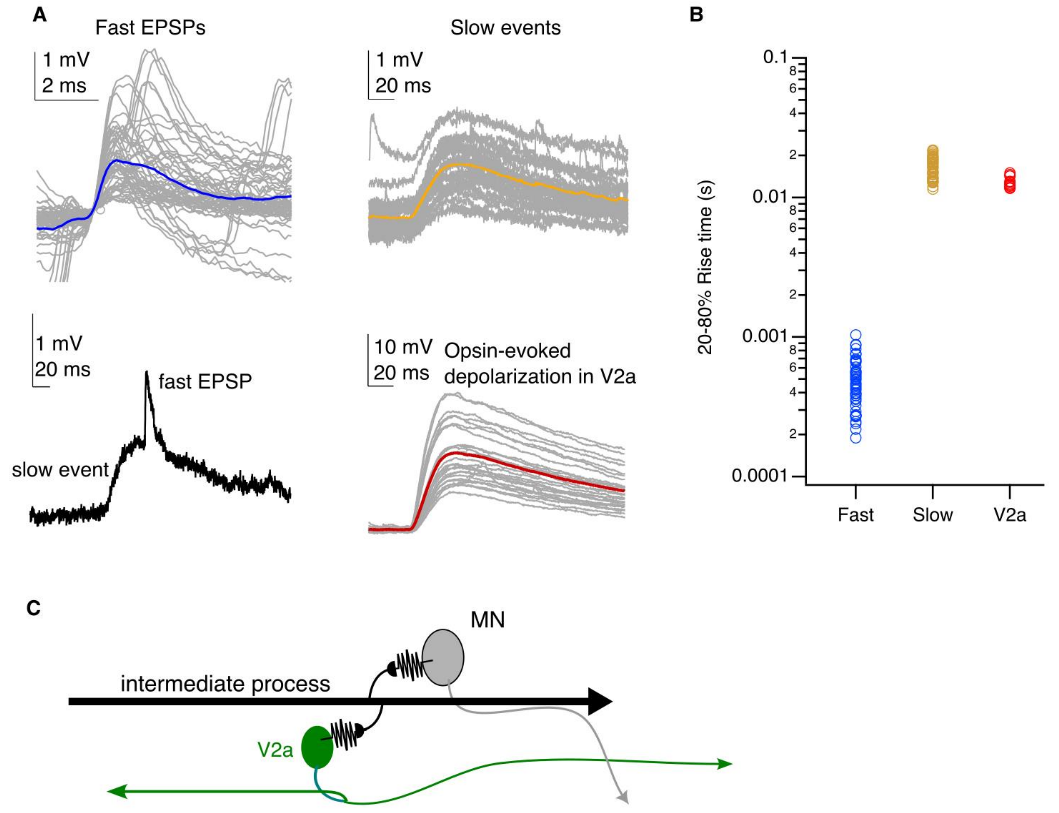


**<u>Supp. Figure S2</u>** Kinetics of events evoked by optical stimulation. Synaptic responses were measured in current clamp for easier comparison with the membrane time constant of V2a neurons. **A,** Top left: Example fast EPSPs recorded in a motor neuron during optical stimulation of V2a neurons located five segments rostral to the recording site. EPSPs are aligned to the middle of the rise (gray) and an average is shown in blue. Top right: Same but for slow events recorded during optical stimulation of V2a neurons in the same segment as the recording site. Note the difference in time scale. Bottom left: A single example trace showing both an underlying slow event and a fast EPSP for comparison. Bottom right: Membrane potential dynamics in a V2a neuron during subthreshold optical stimulation of V2a neurons in that segment. **B,** Summary of rise times for fast EPSPs, slow events, and V2a membrane potential in response to optical stimulation. Rise times of slow events are ~35x longer than fast EPSPs measured in the same neurons. Rise times of V2a subthreshold depolarizations are ~27x longer than fast EPSPs. **C,** These data support a model in which slow events do not report monosynaptic connectivity, either glutamatergic or electrical. Instead, slow events likely represent polysynaptic electrical coupling, with V2a neurons electrically coupled to an intermediate element (local neuron or axon of passage) which is also electrically coupled to the motor neuron, as schematized. Because gap junctions can transmit bidirectionally, in this arrangement V2a depolarization will produce a slow event in the motor neuron whose temporal properties mimic or are slower than the depolarizing event in the V2a neuron, depending on the membrane time constant of the intermediate element.

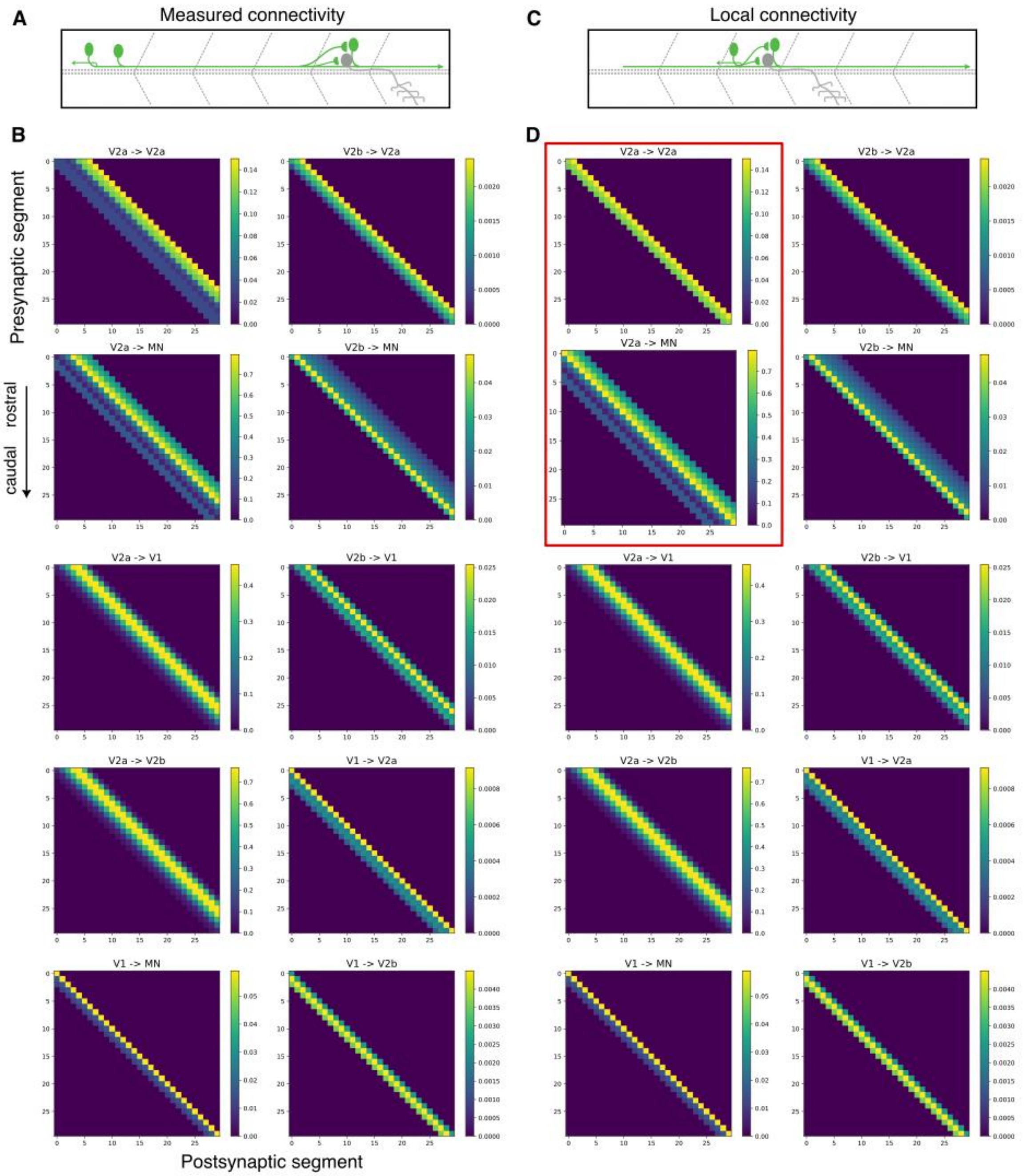


**E**

| population | EL [mV] | C [pF] | gL [nS] | tau_w [ms] | a [nS] | b [pA] | Vr [mV] | VT [mV] | ΔT [mV] |
|---|---|---|---|---|---|---|---|---|---|
| V2a | -60.0 | 13.7 | 1.3 | 50.0 | 0.01 | 5.0 | -42 | -40 | 3.5 |
| MN | -74.0 | 30.0 | 12.5 | 110.0 | 2.2 | 3.0 | -62 | -50 | 1.2 |
| V1 | -76.5 | 23.0 | 3.7 | 5.0 | 4.0 | 50.0 | -55 | -52 | 5.0 |
| V2b | -74.3 | 22.0 | 4.4 | 300.0 | 0.2 | 14.0 | -50 | -45.5 | 3.0 |

**<u>Supp. Figure S3</u>**: Complete network model connectivity and parameters. **A,** Schematic of experimentally measured connectivity. **B,** Connectivity grid between each population of neurons across the 30 segments of the experimentally inspired model. Each segment contained one neuron of each class. **C,** Schematic of local recurrent connectivity. **D,** Connectivity grid between each population of neurons in this recurrent model. Only the two grids outlined by the red box are different between the models. **E,** Cellular parameters of each population, identical between the two models.

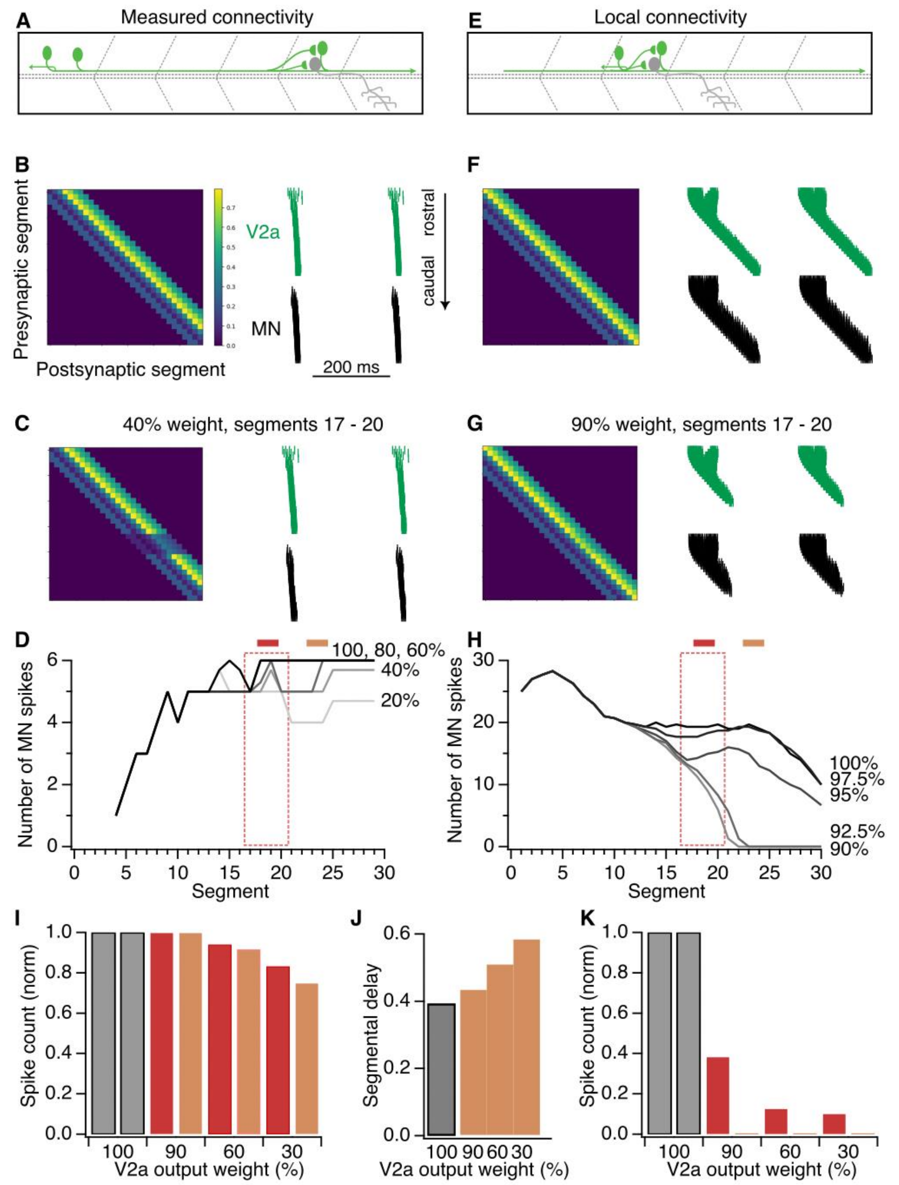


**Supp. Figure S4:** Ablations in mid-caudal region of the network model. As in Fig. 5, but with ablation of synaptic outputs from V2a neurons located in segments 17-20.

**Supplemental Table S1**

Statistical $p$-values for evoked EPSC amplitude comparison with baseline spontaneous activity. To correct for multiple comparisons, values below $p = 0.0038$ (0.05 / 13 segment tests) were considered significant (red).

| | Target neuron class | | | | | |
|---|---|---|---|---|---|---|
| **Segment** | **pMN** | **sMN** | **V2a** | **V1** | **V0v** | **dI6** |
| **6R** | 1.22E-04 | 1.90E-04 | 4.88E-04 | 7.42E-02 | 1.93E-01 | 6.71E-03 |
| **5R** | 1.53E-05 | 1.97E-05 | 1.83E-04 | 1.22E-03 | 2.40E-01 | 7.10E-03 |
| **4R** | 7.63E-06 | 2.50E-07 | 1.53E-05 | 1.16E-03 | 3.98E-02 | 1.00E-02 |
| **3R** | 3.81E-06 | 1.16E-06 | 3.36E-03 | 3.66E-04 | 8.93E-01 | 1.36E-02 |
| **2R** | 2.67E-05 | 1.58E-04 | 1.59E-02 | 6.71E-03 | 5.22E-02 | 1.92E-02 |
| **1R** | 4.22E-03 | 3.65E-04 | 4.20E-04 | 1.03E-02 | 3.95E-01 | 5.69E-02 |
| **0** | 9.54E-06 | 1.15E-01 | N/A | 8.36E-03 | 1.19E-01 | 1.28E-02 |
| **1C** | 1.75E-01 | 5.07E-03 | 3.81E-05 | 7.87E-01 | 6.95E-01 | 1.75E-01 |
| **2C** | 2.63E-01 | 2.50E-04 | 1.00E+00 | 1.00E+00 | 2.32E-01 | 1.05E-01 |
| **3C** | 7.85E-02 | 8.60E-01 | 1.35E-01 | 2.73E-01 | 2.73E-02 | 8.21E-01 |
| **4C** | 3.22E-02 | 8.26E-02 | 7.35E-01 | 6.11E-01 | 3.91E-03 | 9.06E-02 |
| **5C** | 1.56E-02 | 5.42E-01 | 5.57E-01 | 6.85E-01 | 5.47E-01 | 1.35E-01 |
| **6C** | 4.38E-01 | 4.13E-01 | 5.78E-01 | 7.30E-01 | 8.13E-01 | 1.73E-01 |
